# Heavy-tailed distributions in a stochastic gene autoregulation model

**DOI:** 10.1101/2021.06.02.446860

**Authors:** Pavol Bokes

**Affiliations:** Department of Applied Mathematics and Statistics, Comenius University, Bratislava 84248, Slovakia; Mathematical Institute, Slovak Academy of Sciences, Bratislava 81473, Slovakia

**Keywords:** gene expression and regulation, large deviations

## Abstract

Synthesis of gene products in bursts of multiple molecular copies is an important source of gene expression variability. This paper studies large deviations in a Markovian drift–jump process that combines exponentially distributed bursts with deterministic degradation. Large deviations occur as a cumulative effect of many bursts (as in diffusion) or, if the model includes negative feedback in burst size, in a single big jump. The latter possibility requires a modification in the WKB solution in the tail region. The main result of the paper is the construction, via a modified WKB scheme, of matched asymptotic approximations to the stationary distribution of the drift–jump process. The stationary distribution possesses a heavier tail than predicted by a routine application of the scheme.

**MSC 2020:** 92C40; 60J76, 45D05, 41A60

## 1 Introduction

Bursty production of gene products (mRNA or protein molecules) makes an important contribution to the overall gene expression noise [1–4]. Bursts can be modelled as instantaneous jumps of a random process. Burst sizes have been suggested to follow geometric (in a discrete process) or exponential (in a continuous process) distributions [5, 6]; we focus on the latter. Production of gene products is balanced by their degradation and/or dilution. Combining randomly timed and sized production bursts with deterministic decay leads to a Markovian drift–jump model of gene expression [7–10]. More fine-grained models of gene expression are based on a purely discrete [11–15] or a hybrid discrete–continuous state space [16–20]. The drift–jump model can be derived from the fine-grained processes using formal limit procedures [21–26].

In its basic formulation, the drift–jump model for gene expression admits a gamma stationary distribution [27]. The model possesses an explicit stationary distribution also in the presence of a Hill-type feedback in burst frequency [28]. Such regulation can result from common transcriptional control mechanisms [29]. In addition to feedback in burst frequency, there is evidence of feedback mechanisms that act on burst size or protein stability [30–32]. As a specific example of regulation of burst size, the RNA binding protein Puf3 destabilises the mRNA (hence shortening bursts of translation) of COX17 [33]; a synthetic gene encoding for the Puf3 protein while containing untranslated regions of the COX17 gene implements the desired feedback loop [34]. The explicit stationary solution to the drift–jump model has been extended to the case of feedback in protein stability [35]. However, in case of feedback in burst size, an explicit solution is unavailable, save for the special case of Michaelis– Menten-type response [36].

The near-deterministic regime of frequent and small bursts can be analysed using the Wentzel–Kramers–Brillouin (WKB) method; the WKB-approximate solutions closely agree with numerically obtained exact distributions even at moderate noise conditions [37]. Bursty production has been formulated and analysed with the WKB method also in the discrete state space [38–42]. Similar approaches have earlier been used in queueing systems [43, 44]. The standard WKB-type/diffusion-like results are guaranteed to apply for jump-size distributions with super-exponentially decaying tails [45]. Contrastingly, in the sub-exponential case, large deviations are driven by single big jumps [46, 47]. The exponential case can combine both phenomena for random walks: the Cramer/WKB-type result applies in a region of sample space called the Cramer zone, while single big jumps contribute to deviations beyond the Cramer zone [48, 49].

In this paper, the standard WKB-type approach will be shown to be suitable for the drift-jump gene expression model with positive feedback in burst size. If the feedback is negative, the WKB-approach will be shown to apply below a certain threshold (referred to, by analogy with random walks, as the Cramer zone), whereas beyond the threshold (referred to as the tail zone) single big jumps contribute to large deviations. Matched asymptotic approximations to the stationary distribution in the Cramer zone, in the tail zone, and on their boundary will be constructed using a singular perturbation approach [50–53].

The structure of the paper is as follows. Section 2 formulates the model. Section 3 presents the standard WKB approximation scheme. The core of the paper is Section 4, in which the modified WKB scheme is given. The boundary layer is treated in Section 5. The asymptotic results are cross-validated by simulations in Section 6. The paper is concluded in Section 7.

## 2 Model formulation

The drift–jump gene-expression model is a Markov process with piecewise continuous sample paths (Figure 1, left panel). The state *x* of the process represents the concentration of a gene product (say a protein, for concreteness). The discontinuities in the sample path are the production bursts. Between bursts, the protein concentration decays deterministically with rate *γ*(*x*), i.e. as per 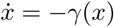. Bursts occur with state-dependent frequency (propensity) *ε*^−1^*α*(*x*). Burst sizes are drawn from an exponential distribution with rate parameter *ε*^−1^*ν*(*x*), in which *x* is the state of the process immediately before the burst; the reciprocal *ε/ν*(*x*) of the rate parameter gives the mean burst size. Decreasing the noise strength *ε* makes bursts more frequent and smaller. The functions *α*(*x*), *ν*(*x*), and *γ*(*x*) can implement feedback in burst frequency, burst size, and protein stability (Figure 1, right panel).

**Figure 1:**
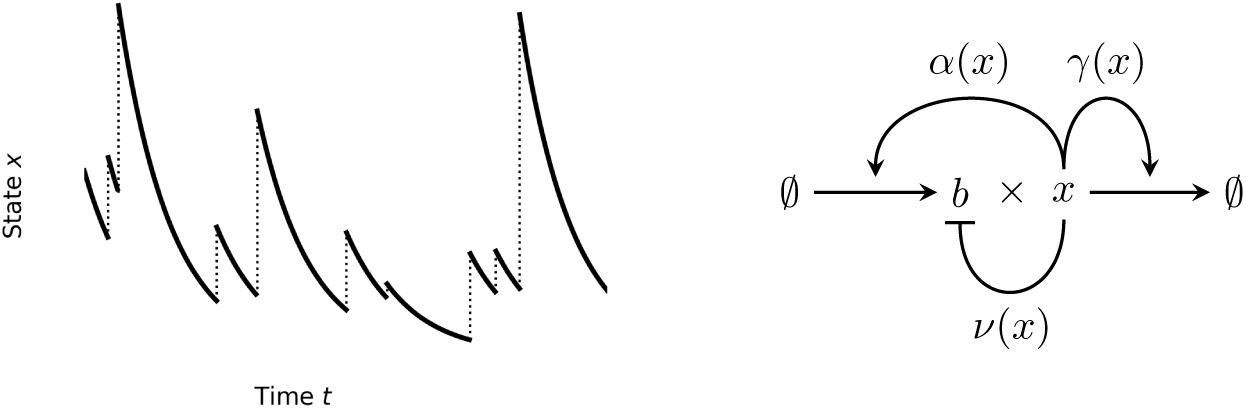
*Left:* Sketch of a typical sample path of the drift–jump gene-expression model. *Right:* Functions *α*(*x*), *ν*(*x*), and *γ*(*x*) quantify feedback in burst frequency, burst size, and protein stability.

The probability density function *p*(*x, t*) of being at state *x* at time *t* satisfies the integro–differential equation

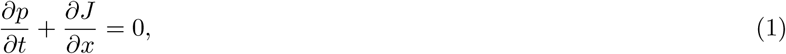

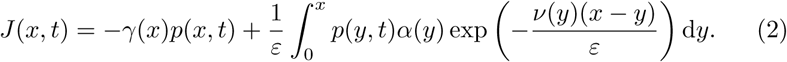

In the conservation equation (1), *J* = *J* (*x, t*) gives the flux of probability across a reference state *x* at time *t*. By (2), it consists of a negative local flux due to deterministic decay and a positive non-local flux due to stochastic bursts. The non-local term integrates, over all states *y < x*, the probability *ε*^−1^*p*(*y, t*) *α* (*y*) that a burst occurs multiplied by the exponential probability that the burst goes beyond the reference state *x*.

Estimating the integral in (2) by the Laplace method [54] as ε → 0, we obtain *J* ∼ (*α*(*x*)*/ν*(*x*) − *γ*(*x*))*p*(*x, t*), which is the probability flux of a purely deterministic process

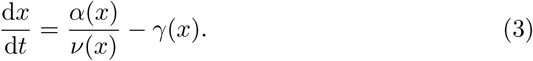

Equation (3) is the deterministic limit of (1)–(2) (sometimes also referred to as the fluid limit or the law-of-large-numbers limit). Retaining a further term in the asymptotic expansion of the non-local term leads to an ad-hoc drift–diffusion approximation to the drift–jump process [55]. Such truncations exhibit different *ε* → 0 asymptotics than the original problem [56].

Equating the flux in (2) to zero, we obtain a Volterra integral master equation

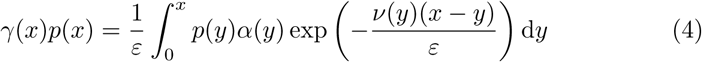

for the stationary distribution. If *ν*(*x*) is constant (no feedback in burst size), then the integral kernel in (4) is both separable and convolution-type [57], and the equation can be solved by differentiation [35] or the Laplace transform [7]. If *ν*(*x*) is non-constant, the kernel is neither separable nor convolution-type, and a general solution seems to be unavailable.

Multiplying a solution *p*(*x*) to (4) by a constant gives another solution. Below, we derive asymptotic approximations to a solution with an arbitrary choice of the multiplication constant, which does not necessarily integrate to one. Normalisation is performed before these approximations are cross-validated by kinetic Monte-Carlo simulations in Section 6.

## 3 Standard WKB scheme

We seek an approximate solution to (4) in the WKB form

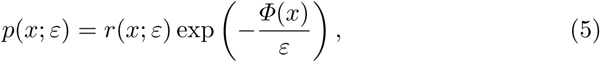

where a regular dependence

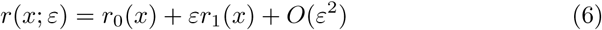

of the prefactor on *ε* is postulated. The function *Φ*(*x*) in (5) is referred to as the (quasi)potential. The ansatz (5)–(6) is equivalent to the frequently encountered alternative form *p*(*x*; *ε*) = exp(−*ε*^−1^(*Φ*_0_(*x*)+*εΦ*_1_(*x*)+*ε*^2^*Φ*_2_(*x*)+…)) of the WKB expansion [58]; the correspondence between the terms is given by *Φ*(*x*) = *Φ*_0_(*x*), *r*_0_(*x*) = exp(−*Φ*_1_(*x*)), and *r*_1_(*x*) = −exp(−*Φ*_1_(*x*))*Φ*_2_(*x*).

Inserting (5) into (4) gives

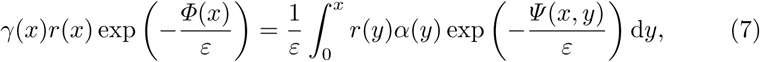

where

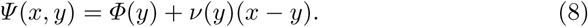

Differentiating (8) with respect to *y* and setting *y* = *x* gives the relations

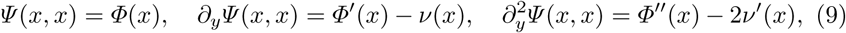

which tie up the local behaviour of *Ψ* (*x, y*) near the boundary *y* = *x* and that of the (yet unknown) potential.

Provided that

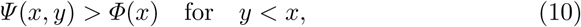

the dominant contribution to the integral on the right-hand side of (7) comes from an *O*(*ε*)-wide neighbourhood of the right boundary. Estimating the integral in (7) by the Laplace method and cancelling the common exponential term gives

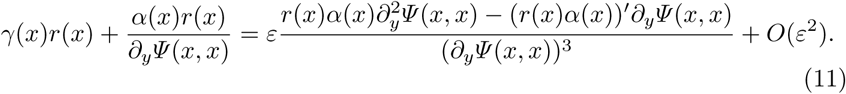

Inserting (6) and (9) into (11), and collecting *O*(1) terms, yields the potential

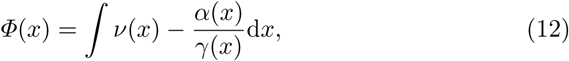

while collecting *O*(*ε*) terms determines the prefactor (Appendix A)

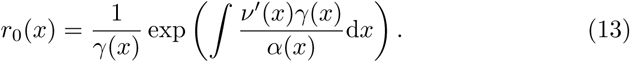

The constants of integration in the indefinite integrals in (12)–(13) add up to the normalisation constant in the probability distribution (5) and can be chosen arbitrarily.

The weak point of this section is the assumption (10). Combining (8) and (12), we see that

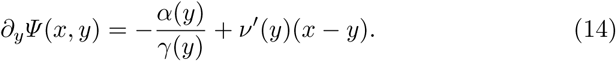

If *ν*(*x*) is decreasing (positive feedback case), then ∂_*y*_*Ψ* (*x, y*) *<* 0 for *y* ≤ *x*, which confirms (10) post hoc. If *ν*(*x*) is constant (no feedback in burst size), then *r*_0_(*x*) exp(−*Φ*(*x*)*/ε*) given by (12)–(13) is the exact solution to (4) [35]. The case of an increasing *ν*(*x*) (negative feedback in burst size) is the subject of the rest of the paper; it requires a modification in the WKB scheme.

## 4 Modified WKB scheme

Since the derivation of (12) involved a local estimate of the integral in the Volterra master equation (7), we refer to the function *Φ*(*x*) as the local potential. We assume that it satisfies the conditions

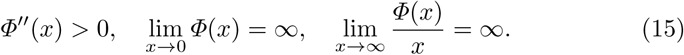

An example of a local potential which satisfies (15) is given by the parametric choice

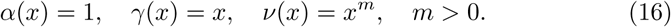

The constancy of the burst frequency and the linearity of the decay rate in (16) means that feedback occurs only in the burst size. The coefficient *m* can be interpreted as the number of protein molecules that need to cooperate to repress a production burst. Although the analysis of this section is performed for a general parametric choice, all graphical examples pertain to the choice (16). Specialised calculations for (16) can be found in Appendix B.

We define the modified potential by

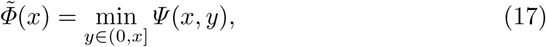

where *Ψ* (*x, y*) is given by (8) and (12). Since *Ψ* (*x, x*) = *Φ*(*x*), we immediately obtain 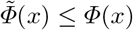. Graphical examination shows that there is a critical point *x*_*_ such that 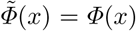 for *x* ≤ *x*_*_, whereas for *x* ≥ *x*_*_ the graph of 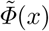 is the envelope of rays *x* → *Ψ* (*x, y*) parametrised by *y* (Figure 2, left). The region *x < x*_*_ will be referred to as the Cramer zone, and the region *x > x*_*_ as the tail zone.

**Figure 2:**
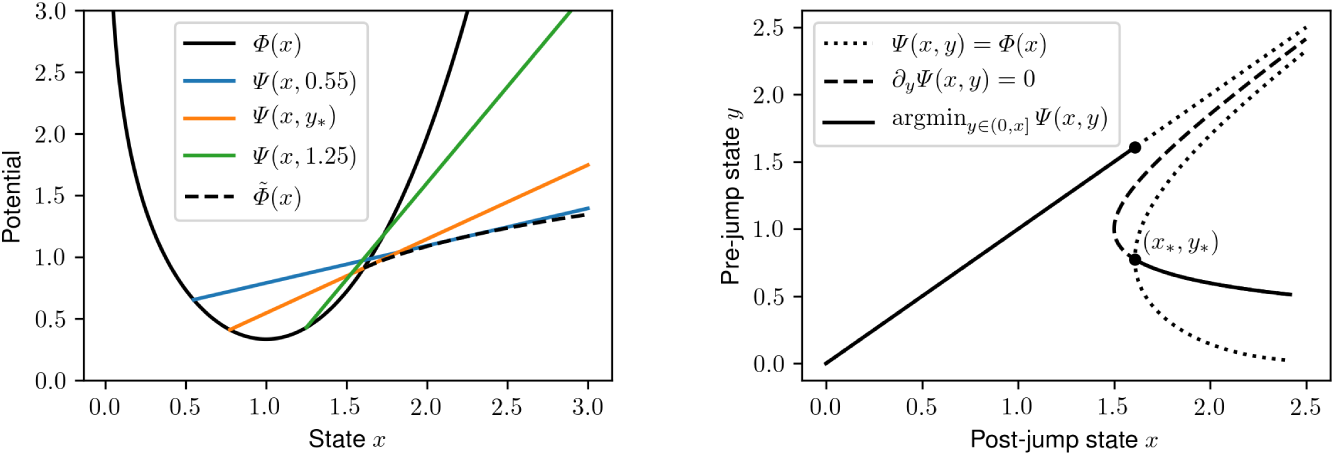
*Left:* The local potential *Φ*(*x*) and the modified potential 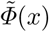 as the envelope of rays *Ψ* (*x, y*). *Right:* Important curves in the domain of *Ψ* (*x, y*). The parametric choice (16) with *m* = 2 is used.

The critical point *x*_*_ is the supremum of all *x* for which (10) holds. It follows that at *x* = *x*_*_, an internal minimiser *y* = *y*_*_ *< x*_*_ of (17) exists in addition to the boundary minimiser *y* = *x*_*_, so that

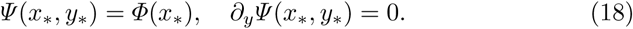

Equations (18) determine the critical pair (*x*_*_, *y*_*_) uniquely (Figure 2, right).

In the tail zone,

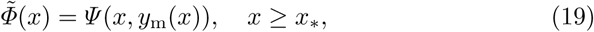

where *y*_m_(*x*) is the internal minimiser, which satisfies

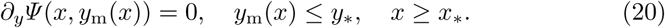

Note that *y*_m_(*x*_*_) = *y*_*_ and *y*′_*m*_ (*x*) *<* 0. The potential derivative satisfies

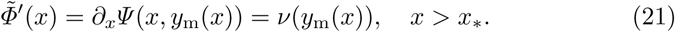

The first equality in (21), which states that the envelope is tangential to the ray that forms it, is a simple example of an envelope theorem [59].

The local potential and the envelope of rays intersect at *x* = *x*_*_ transversally with slopes 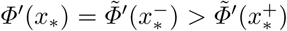: the modified potential has a discontinuous derivative at *x* = *x*_*_. Convexity of *Φ*(*x*) and concavity of 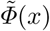 imply 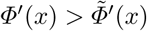 for *x* > *x*\*. These comparisons, together with (12) and (21), lead to the inequality

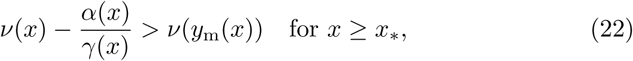

which will be important in the subsequent analysis.

If we look for a solution *p*(*x*; *ε*) to (4) in a form that is logarithmically equivalent to 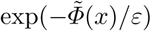, then the integrand on the right-hand side is logarithmically equivalent to 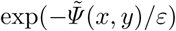, where

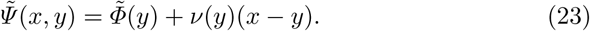

For *x > x*_*_, we have (Appendix C)

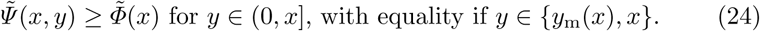

Notably, 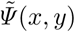 — as function of *y* ∈ (0, *x*] — is minimised both on the right boundary and internally, whereas *Ψ* (*x, y*) is minimised only internally for *x > x*_*_. The integral on the right-hand of (4) side will be logarithmically equivalent to 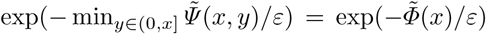, which is the asymptotics postulated for the solution. The use of the modified WKB potential (17) thus leads to a desired balance between the sides, at least to a logarithmic precision, of the master equation (4).

Important contributions to the integral term in (4) come from the neighbourhoods of the minimisers *y* = *y*_m_(*x*) and 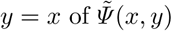 for a fixed *x > x*_*_. The function is locally parabolic near the internal minimiser *y* = *y*_m_(*x*) *< x*_*_, but it is locally linear near the boundary minimiser *y* = *x > x*_*_. By the Laplace method [54], an *O*(*ε*^1*/*2^) neighbourhood of the parabolic minimiser, but only an *O*(*ε*) neighbourhood of the linear minimiser, contribute. The internal minimiser lies in the Cramer zone, where the standard procedure of Section 3 yields

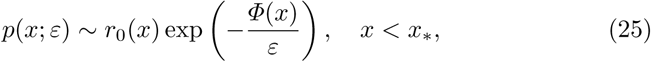

with the prefactor defined by (13). In order to balance the contributions of the internal and boundary minimisers, we compensate at the level of prefactor, seeking the solution outside the Cramer zone in the form of

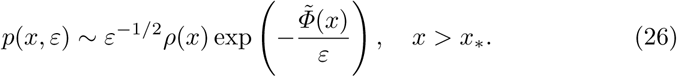

Inserting the WKB expansions (25) and (26) into the Volterra master equation (4), we find that for a *δ* ⟫ *ε*^1*/*2^ we have

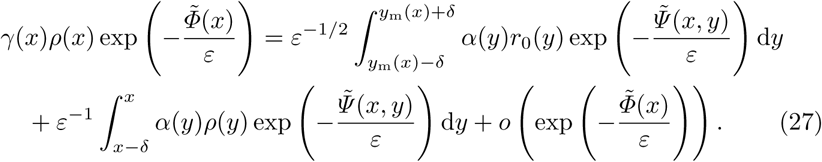

Estimating the integrals by the Laplace method, cancelling the common exponential term, and collecting at the leading order, we obtain

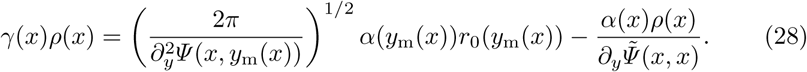

Differentiating (23) with respect to *y* and using (21) gives

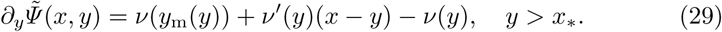

We set *y* = *x* into (29) and insert the result into (28), arriving at

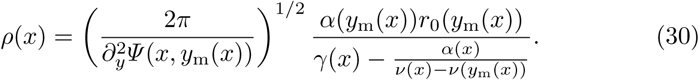

Inequality (22) ensures that the denominator in (30) is positive.

In the next section, we complete the approximation scheme by constructing an inner solution in a neighbourhood of the Cramer boundary *x* = *x*_*_ that matches (25) to the left and (26) to the right.

## 5 Boundary layer

The discontinuity in the potential derivative and the mismatch of prefactor magnitudes in (25) and (26) suggest the presence of a boundary layer near *x* = *x*_*_. We define the inner variable *ξ* via the transformation

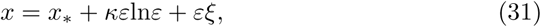

where the constant *κ >* 0 will be specified later. Qualitatively, as *x* increases towards *x*_*_, the integral in the Volterra equation (7) begins to feel the “ghost” of the internal minimum of *Ψ* (*x*_*_, *y*) at *y* = *y*_*_; the local approximation scheme of Section 3 breaks down before *x*_*_ is reached.

The inner solution is sought to be proportional to a regular function of the inner variable:

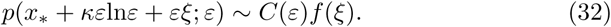

Inserting (32) into the Volterra master equation (4) and seeking a dominant balance between terms gives (Appendix D)

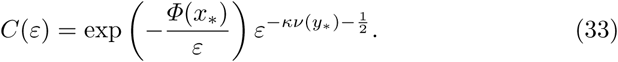

Collecting the leading-order terms in the expansion of (4), the unknown *f* (*ξ*) is found to satisfy an inhomogeneous Volterra equation (Appendix D). Unlike the original equation (4) for *p*(*x*; *ε*), the Volterra equation for *f* (*ξ*) has a separable kernel, and is readily turned into a differential equation with a general solution (Appendix D)

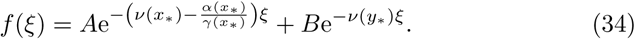

The constant

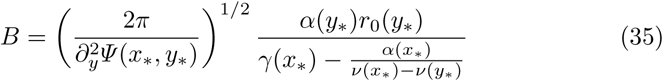

multiplying the particular solution in (34) is found by the method of undetermined coefficients; the constant of integration *A* multiplying the homogeneous solution remains undetermined at this stage.

The *ξ* → ∞ asymptotics of (32) agree with the *x* → *x*_*_ behaviour of the tail-zone WKB solution (26) with arbitrary choices of the integration constant *A* and the constant *κ* in the offset of the boundary layer (31) (Appendix E). In order to determine the two constants, the *ξ* → −∞ asymptotics of (32) need to be matched to the *x* → *x*_*_ behaviour of the Cramer-zone WKB solution (25). By inequality (22) with *y*_m_(*x*_*_) = *y*_*_, the first term in (34) dominates as *ξ* → −∞, so that

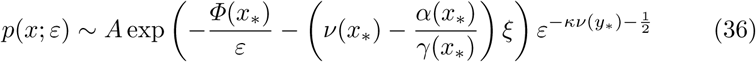

in the overlap of the inner solution and the outer (Cramer-zone WKB) solution. On the other hand, inserting (31) into the outer solution (25) gives

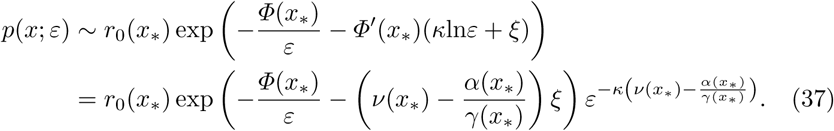

Comparing (36) to (37) yields

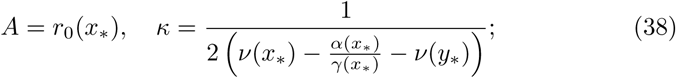

inequality (22) thereby guarantees that *κ >* 0 as advertised at the beginning of the boundary-layer analysis. Equations (38) complete the inner solution and thus the asymptotic analysis of (4).

## 6 Numerical solution

Before being compared to a numerical solution, the asymptotic solutions are normalised by

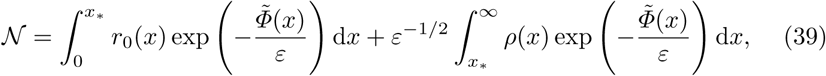

which is calculated by numerical quadrature.

For the numerical solution, sample paths *x*_*i*_(*t*), *i* = 1, …, *N*, 0 ≤ *t* ≤ *T*, subject to *x*(0) = *x*_0_ are generated using the exact stochastic simulation algorithm (Appendix F). The solution is constructed by the histogram method from the dataset of final-time values {*x*_*i*_(*T*)}_*i*=1,…,*N*_. Specifically, we divide an interval [0, *x*_max_] into *n* equally sized bins, count the number of data in each bin, and divide the counts by *Nx*_max_*/n* so as to normalise into a probability density. The histogram estimate is close to the exact solution *p*(*x*; *ε*) to the Volterra master equation (4) if the number of samples *N* is large (so that the statistical error is small) and the simulation end time *T* is large (so that the process equilibrates to steady state).

Figure 3 compares the three matched asymptotic approximations to the numerical solution for selected values of the noise strength *ε*. Decreasing *ε* leads to a close agreement between the numerical solution and the asymptotic approximations in their respective regions of validity (Figure 3, top panels). As *ε* decreases further (Figure 3, bottom panels), the Cramer-boundary and tail behaviour become exponentially improbable, and cannot be reliably estimated from a feasible number (say a billion) of samples. Nevertheless, the chosen examples demonstrate that the naive solution, which extends (25) outside the Cramer zone, underestimates the tail of the stationary distribution, whereas the alternative approximations provide an adequate description.

**Figure 3:**
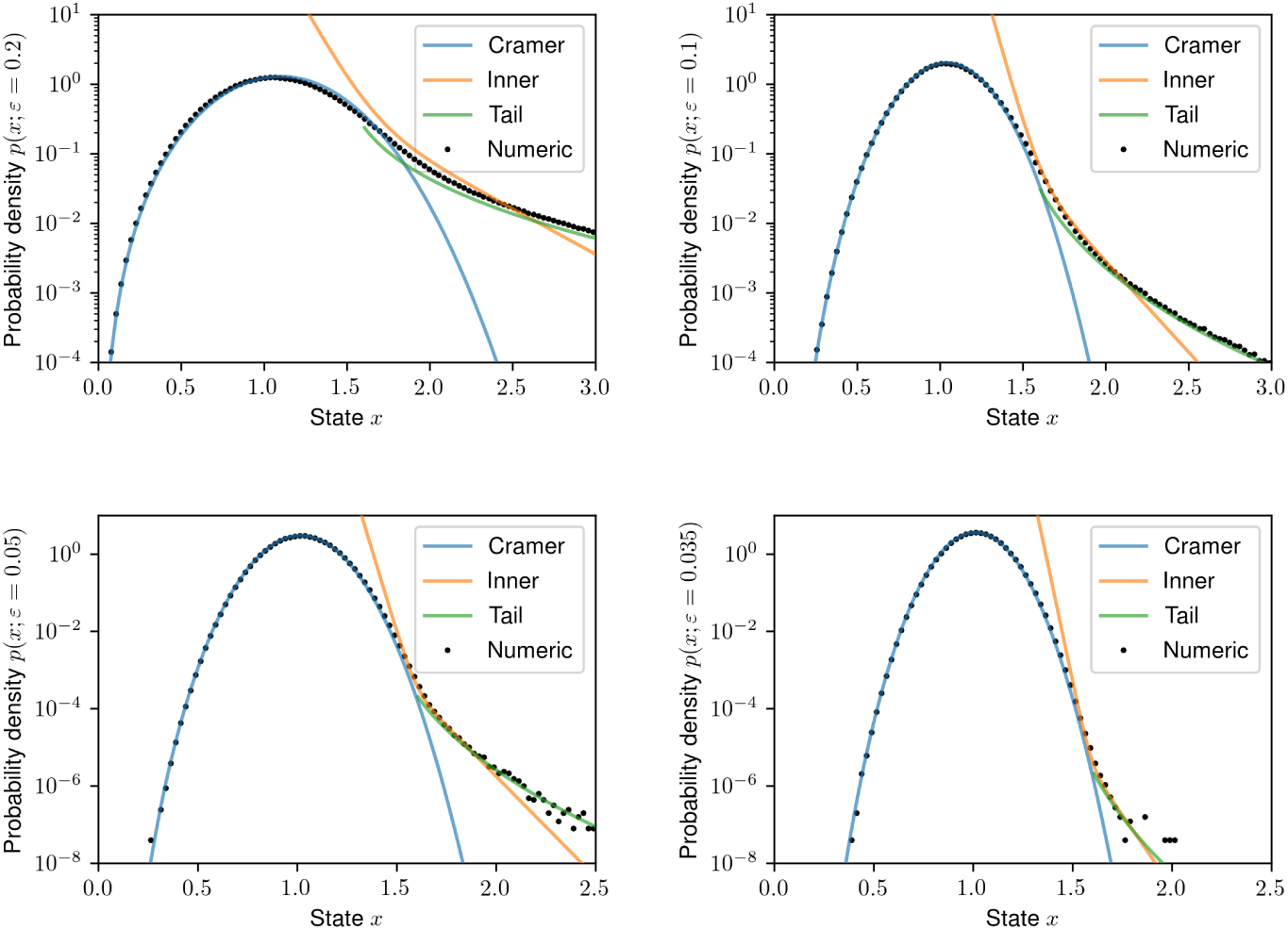
The simulation-based probability density (dots) is compared to the individual asymptotic approximations (solid lines), namely the WKB solution in the Cramer zone (25), the inner solution in the boundary layer (34), and the WKB solution in the tail zone (26). *Model parameters:* we use (16) with *m* = 2; values of *ε* are specified in the label of the ordinate. *Numerical parameters: x*_0_ = 1, *T* = 30, *N* = 10^8^ (upper panels), *N* = 10^9^ (lower panels), *n* = 100, *x*_max_ = 3 (upper panels) and *x*_max_ = 2.5 (lower panels).

## 7 Conclusion

This paper provides matched asymptotic approximations to the stationary distribution of a drift–jump model for stochastic gene expression. The analysis revolves around the estimation of the integral term in the Volterra master equation (4). The integral term represents the flux of probability due to production bursts through a reference state *x*. In the Cramer region (*x < x*_*_), the flux consists solely from local contributions (*y* ≈ *x*), whereas in the tail region (*x > x*_*_), a contribution comes also from within the interval. The latter corresponds to the ‘single big jumps’ advertised in the abstract.

Negative feedback in burst size is a prerequisite for the singular behaviour in question. Conceptually, in the presence of negative feedback in burst size, it is ‘cheaper’ to hunker down and then take a giant leap, than to climb up with tiny steps. The result is thus in agreement with the broad principle that any large deviation occurs in the least unlikely of all the unlikely ways [60].

For a power-law feedback in burst size (16), an earlier study [61] provides the asymptotic formulae *p*(*x*) ∼ *x*^−1+1*/ε*^ as *x* ⟶ 0 and *p*(*x*) ∼ *ηx*^−1−1*/εm*^ as *x* → ∞ (for an *ε >* 0 which is not necessarily small), where *η* = *m*^−1^*ε*^−1+1*/εm*^ Γ (1*/εm*). The latter asymptotic implies that the stationary distribution is heavy-tailed (sub-exponential) [62]. The same study establishes a central-limit-theorem-type approximation that is valid as *ε* → 0 for |*x*−1| = *O*(*ε*^1*/*2^) [61]. The current study contributes by approximations that apply as *ε →* 0 throughout the state space *x >* 0. In particular, elementary calculations show that the *x* → 0 behaviour of the Cramer-zone WKB solution and the *x* → ∞ behaviour of the tail-zone WKB solution are equal to the aforementioned *x* → 0 and *x* asymptotics of the exact solution, wherein *η* ∼ (2*π/εm*)^1*/*2^e^−1*/εm*^*m*^−1*/εm*^ for *ε* « 1 by Stirling’s formula.

While the power-law non-linearity is the principal example of this paper, the popular Hill function can be reduced to the power non-linearity by an explicit transformation [61]. For feedback responses that do not satisfy the constraints introduced in Section 4, one expects that multiple disconnected Cramer zones may exist, in which the potential is evolved locally, and which are interspersed by zones in which the potential is formed by an envelope of rays.

It is also expected that the current methods can be applied to the discrete framework, if this is extended so as to include feedback in burst size, with burst sizes drawn from the geometric distribution [5]. More widely, the current results can be pertinent to other fields, e.g. to the Takács equation for the amount of unfinished work in an M/M/1 queue, or other jump processes with jump measures with exponential tails [44].

Earlier studies argue that the subtleties that arise with feedback in burst size are an artefact of delay [36, 37]. Indeed, the memoryless property of the exponential distribution of burst sizes implies a lack of control at the infinitesimal timescale of burst growth. In light of this argument, the results contribute to the understanding of the interplay between bursting and delay [63–67].

## Appendix A: Solving the ODE for the prefactor

The prefactor satisfies the ordinary differential equation

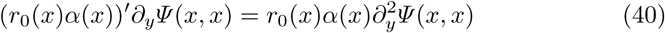

with solution

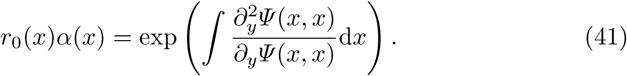

Using (9) and (12), we obtain

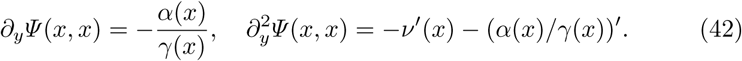

Inserting (42) into (41) gives

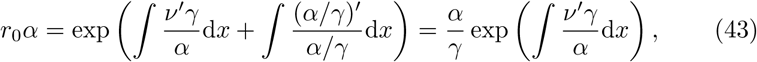

from which (13) follows.

## Appendix B: The power-law example

For the choices (16), the Cramer-zone potential (12) and prefactor (13) are given by

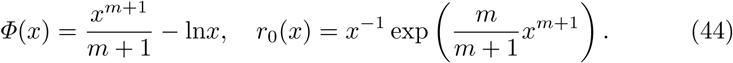

The rays (8), whose envelope constitutes the modified potential, are given by

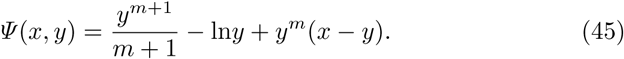

The derivative of the rays with respect to the envelope parameter *y* is given by (14), which is

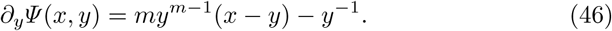

The critical pair (*x*_*_, *y*_*_) is the solution of the non-linear system (18), which here takes the form

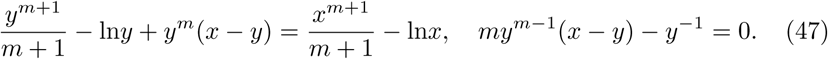

The author solved (47) iteratively, starting from an initial guess *x* = 1 + 1*/m* and *y* = 1, using Broyden’s first Jacobian approximation method [68].

In order to find the internal minimiser *y* = *y*_m_(*x*) of *Ψ* (*x, y*), we are required by (20) to solve for a given *x* ≥ *x*_*_ the equation

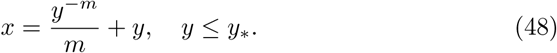

The author did this in two steps: first, he found *x*_*i*_ corresponding to the values *y*_*i*_ = *y*_*_(1 − *i/I*), *i* = 0, 1, …, *I* − 1, by substitution into (48); second, he used a cubic spline to interpolate *y* = *y*_m_(*x*) between *y*_*i*−1_ = *y*_m_(*x*_*i*−1_) and *y*_*i*_ = *y*_m_(*x*_*i*_), *i* = 1, …, *I* − 1. The modified WKB potential and prefactor are calculated by substituting the spline of *y* = *y*_m_(*x*) into (19) and (30).

The formulae (30) and (35) for the tail-zone WKB and boundary-layer solutions evaluate 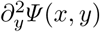 at the internal minimiser *y* = *y*_m_(*x*). Differentiating (46) gives

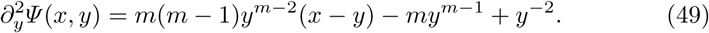

At *y* = *y*_m_(*x*) the first derivative (46) of *Ψ* (*x, y*) vanishes, and the first term in (49) simplifies to (*m* − 1)*y*^−2^; therefore,

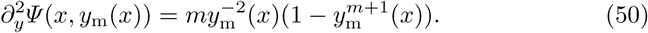

Note that *y*_m_(*x*) *< y*_*_ *<* 1, so that (50) is positive, as required in (30) and (35).

## Appendix C: Minimisers of 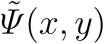

Let us investigate the behaviour of 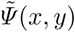 as function of *y* ∈ (0, *x*] for a fixed *x > x*_*_. The Cramer and the tail regions are thereby treated separately:

1. *y* ≤ *x*_*_. Here we have

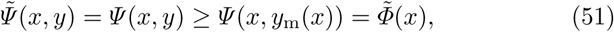

with equality in place if *y* = *y*_m_(*x*).
2. *y* ≥ *x*_*_. Here

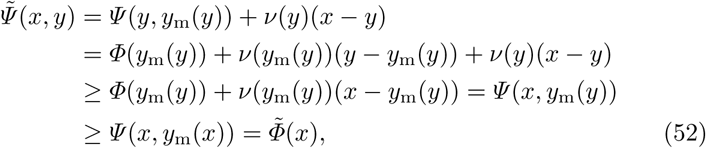

with both estimates becoming equalities if *y* = *x*; the first estimate requires that *ν*(*x*) be non-decreasing (negative feedback in burst size).

## Appendix D: The inner solution

We divide the integration interval in (4) into 0 *< y < x*_o_ and *x*_o_ *< y < x*, where *x*_o_ belongs to the overlap of the WKB approximation (25) and the inner approximation (32).

In the first interval, the integral is estimated by means of the WKB approximation (25) and the Laplace method as

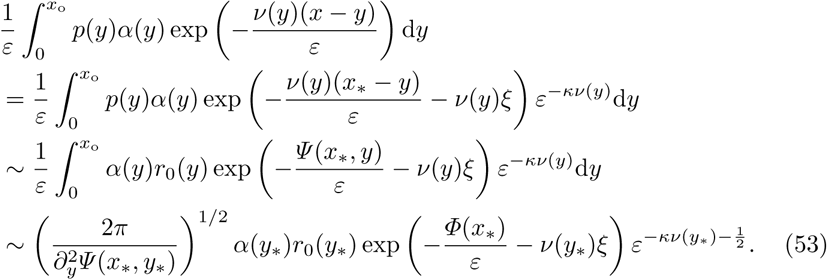

In the second interval, the substitution *y* = *x*_*_ + *κε*ln*ε* + *εη* and the inner approximation (32) give an asymptotic estimate

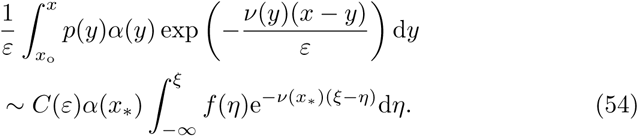

Requiring that (53) and (54) be of the same order implies (33) for the proportionality constant *C*(*ε*) in the inner solution (32).

Inserting (32), (53), and (54) into the Volterra master equation (4), and then dividing by *C*(*ε*), yields an inhomogeneous Volterra equation for *f* (*ξ*) with a separable kernel:

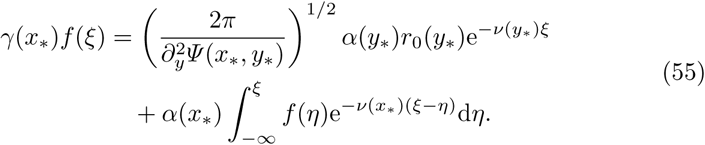

Multiplying (55) by e^*ν*(*x*^*)*ξ* and differentiating with respect to *ξ* turns the integral equation (55) into a differential equation

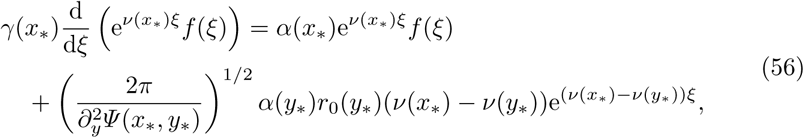

which has a general solution (34)–(35).

## Appendix E: Matching to the right

By inequality (22), the second term in the inner solution (34) dominates for *ξ* → ∞; inserting it and (33) into (32) gives

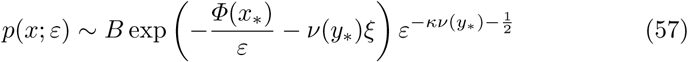

in the overlap of the inner solution and the outer solution to its right.

On the other hand, inserting the transformation (31) into the outer solution (26), re-expanding, and using (21) gives

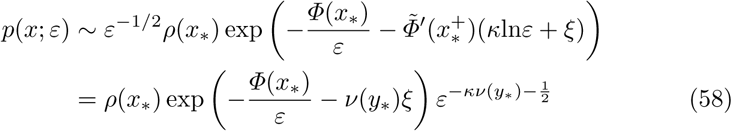

in the overlap. Comparing (57) and (58), we find *B* = *ρ*(*x*_*_), which is consistent with (30) and (35).

## Appendix F: Stochastic simulation algorithm

Here we provide a stochastic simulation algorithm that can be used to generate a sample path *x*(*t*) of the process on a time interval [0, *T*] subject to an initial condition *x*(0) = *x*_0_. Similarly like the well-known Gibson–Bruck/Gillespie algorithm, the algorithm does not introduce truncation errors, but only statistical and round-off errors, and in this specific sense it is an exact simulation algorithm. For simplicity, we focus on the situation when the feedback acts only on burst size but not on burst frequency or protein stability; the general case is discussed in the end of the appendix.

Each sample path is generated iteratively as follows. Assume that the sample path *x*(*t*) has already been generated on an interval 0 ≤ *t* ≤ *t*_cur_ (initially *t*_cur_ = 0 and *x*(0) = *x*_0_ is an initial value). Assuming the absence of feedback in burst frequency (*α*(*x*) = 1), the exponentially distributed waiting time until the coming burst is sampled by the inversion method as

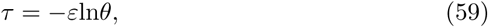

where *θ* is drawn from the uniform distribution in the unit interval. Assuming the absence of feedback in protein stability (*γ*(*x*) = *x*), the sample path decays exponentially until the coming burst:

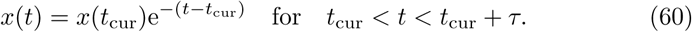

At the time of the next burst the sample path is increased by the exponentially distributed burst size:

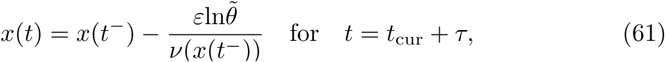

where *x*(*t*^−^) = *x*(*t*_cur_)e^−*τ*^ denotes the state of the sample path immediately before the burst; the variate 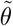 is drawn from the uniform distribution in the unit interval independently of *θ*. Thus one round of iteration via (59), (60), and (61) extends the sample path from the interval [0, *t*_cur_] to the interval [0, *t*_cur_ +*τ*]. The algorithm is repeated until the state *x*(*T*) at a required end time *T >* 0 is found.

The algorithm can be modified to account for feedback in burst frequency and protein stability. If feedback in burst frequency is present, the waiting time needs to be drawn from a distribution with a non-constant hazard function [8]. If feedback in protein stability is present, the sample path needs to be evolved as per 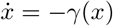 between bursts.

